# Vestibular signals enhance the temporal precision of predictive movements to musical rhythms

**DOI:** 10.64898/2026.09.24.754009

**Authors:** Ryoichiro Yamazaki, Junichi Ushiyama

## Abstract

Humans exhibit head movements synchronized with musical rhythms, frequently observed during instrument playing or dancing (e.g., hip hop). We conjectured that vestibular input from head movements could aid motor execution anticipatory of musical rhythms, known as sensorimotor synchronization (SMS). To examine this, we applied subthreshold galvanic vestibular stimulation (GVS) simultaneously with musical beats while healthy adults performed SMS to those beats. We designed a GVS waveform where the current rose steeply before each beat onset, which was expected to trigger intense vestibular afferent firing before the beats (as actual head movements would). Participants flexed their index finger in synchrony with the beats of syncopated auditory rhythms. In GVS trials, the SMS timing variability was significantly lower than in trials without GVS, suggesting that vestibular input stabilizes SMS to musical beats. Additionally, a control experiment using a waveform with a gradually increasing current revealed no significant difference in SMS variability between trials with and without GVS. This study provides causal evidence that the vestibular system is involved in synchronized movement to music. In addition, the results suggest that a specific parameter of the GVS waveform could influence predictive motor execution anticipatory of upcoming auditory events.

## Introduction

Head movements synchronized to musical rhythms are often seen during musical instrument playing and dancing such as hip hop (Burger, Thompson, Luck, Saarikallio, & Toiviainen, 2013; Palmer, Spidle, Koopmans, & Schubert, 2019; Swarbrick et al., 2019; Van Dyck et al., 2013). Such periodic head movements are involved in the perception of musical rhythm. Palmer et al. (2019) reported that singers exhibited increased head movements in time with the beat when synchronizing to the beat was difficult during duets, suggesting that head movements may contribute to beat synchronization. Phillips-Silver and Trainor (2005, 2007) reported that the periodicity of head movements experienced when listening to music biases the perception of meter in subsequently presented auditory rhythms. This phenomenon has been attributed to vestibular inputs (Trainor, Gao, Lei, Lehtovaara, & Harris, 2009). Studies investigating the influence of vestibular input on the perception of musical rhythm have primarily dealt with training effects (Phillips-Silver & Trainor, 2005, 2007, 2008; Tichko, Kim, & Large, 2021; Tichko & Large, 2019; Trainor et al., 2009). However, we suspected that a mechanism that provides immediate gains for motor performance also exists, underlying the habit of moving one’s head to ongoing music.

We have previously demonstrated that, in a sensorimotor synchronization (SMS) task requiring index finger flexion to a musical beat, trials where the head was moved simultaneously with the beat exhibited higher timing stability during SMS compared with trials without head movements (Yamazaki & Ushiyama, 2024). This finding suggests that voluntary head movement plays a positive role in real-time auditory rhythm processing and SMS to that rhythm. However, because head movements generate various sensory inputs, this result does not provide direct evidence that the effect arises from vestibular input induced by head movements. Addressing this point would advance our essential understanding of head movements to music rhythms, which are shared across cultures and species (Fitch, 2013; Palmer et al., 2019; Schachner, Brady, Pepperberg, & Hauser, 2009; Swarbrick et al., 2019; Van Dyck et al., 2013). Furthermore, rhythm perception and the generation of motor rhythms are integral to daily activities like speech and gait. Therefore, elucidating the role of vestibular input in these functions is crucial for comprehension of the neural bases underpinning human behavior in daily life.

Here, using a technique to electrically stimulate the vestibular system [galvanic vestibular stimulation (GVS)], we examined the causal effect of vestibular input on synchronized movement to musical beats. If SMS performance is enhanced only when GVS is applied, as in our previous head movement experiments, this would support the hypothesis that vestibular input accompanying head movement enhances predictive motor execution anticipatory of musical rhythm. In the experiments, instantaneous sub-threshold GVS was applied synchronously with the beat in half of the trials to mimic the situation where the head moves to the beat. SMS performance was compared between trials with GVS (GVS condition) and without GVS (control condition) to examine the immediate effect of vestibular input on SMS. We conducted two experiments using different GVS waveforms with varying parameter settings, which was expected to affect the firing rates of vestibular afferent fibers differently. The experiment using GVS with a steep increase in current showed improved SMS timing stability in the GVS condition compared with the control condition, whereas the experiment using GVS with a gradual increase in current did not. These experiments revealed that vestibular input causally modulates SMS performance and suggested that the slope of the current could be a potential determinant of this effect.

## Methods

### Participants

In total, 36 (10 women and 26 men; mean age, 21.03 ± 1.84 years) and 23 (11 women and 12 men; mean age, 21.43 ± 2.66 years) healthy young adults were recruited to experiments 1 and 2, respectively. The sample size was determined with G*Power 3.1.9.4 (Faul, Erdfelder, Lang, & Buchner, 2007), assuming a two-tailed one-sample t-test with an effect size of 0.6, statistical power of 0.8, and an alpha level of 0.05. The following participants were excluded from the analyses: (1) those with a history of neurological disease or impaired auditory or motor function and (2) those unable to correctly synchronize their movements to the stimuli in practice or experimental sessions. Exclusion on the basis of incorrect synchronization depended on the consistency between the numbers of movements and beats (i.e., 64): those who showed a larger or smaller number of finger flexions versus the number of beats were excluded. Ultimately, we analyzed 20 participants in experiment 1 and 20 in experiment 2. At the beginning of each experiment, we used the Japanese version of the Flinders Handedness survey (Nicholls, Thomas, Loetscher, & Grimshaw, 2013; Okubo, Suzuki, & Nicholls, 2014) to assess handedness. All participants were right-handed; their handedness scores ranged from 8 to 10 in experiment 1 and from 9 to 10 in experiment 2.

All experimental procedures were conducted in accordance with the Declaration of Helsinki (except the step of pre-registration in a database) and approved by the Research Ethics Committee at Shonan Fujisawa Campus, Keio University (approval number 515). GVS was in compliance with a published assessment confirming minor adverse effects of GVS (Utz et al., 2011). The participants received sufficient explanations about the purpose and methods of the experiments and provided written informed consent in advance of participation.

### Auditory Stimuli

A countdown consisting of eight tones was presented, immediately followed by a single rhythm pattern repeated eight times consecutively in each trial. The rhythm patterns were created in the same manner as in Large, Herrera, and Velasco (2015). We adjusted “syncopation intensity” [defined as the number of quarter notes shifted from the on-beat to the off-beat within the eight notes in a 2-bar repetition unit, compared with a 4/4 metronome as the baseline (Level 0)] for each participant. Accordingly, rhythm patterns with only one note shifting from eight evenly spaced quarter notes were defined as the “weakest syncopation intensity” (Level 1), whereas patterns with five notes shifted were defined as the “strongest syncopation intensity” (Level 5) (Supplementary Figure S1). This operation is based on the principle that syncopation intensity increases when a tone is placed on a weak meter structure rather than on a strong meter structure, and a rest is placed on a strong meter structure (Witek, Clarke, Kringelbach, & Vuust, 2014). Syncopation patterns created in this manner induce beat perception at the quarter-note level (Chapin et al., 2010; Tal et al., 2017; Vuust, Heggli, Friston, & Kringelbach, 2022). Level adjustments were made during the practice session (see the *SMS Task* section below). For Level 4, 6 patterns were randomly selected from the 10 patterns used in Chapin et al. (2010). For all other levels, 6 rhythm patterns were randomly generated for each participant to meet the level definition. The rhythm patterns used in the practice and experimental sessions were generated separately, and were controlled so that no patterns were shared between sessions. The countdown and syncopation rhythm were presented with pure tones at 220 and 440 Hz, respectively. The amplitude of each tone rose linearly from 0 to the optimal volume over 10 ms, was sustained for 30 ms, and then fell linearly to 0 over 10 ms. In the experimental session, auditory stimuli were presented in 4/4 time at 100 bpm: the interval between the perceived beats at 4/4 meters was 600 ms (i.e., 1.67 Hz). Meanwhile, in the practice session, either 95 or 105 bpm was randomly assigned equally to each participant, preventing them from learning the tempo for the experimental session in advance. To minimize the impact of environmental noise, we added white noise to each stimulus. The amplitude of the white noise increased linearly from the second countdown, reaching the same amplitude as the tones at the second 4/4 beat of the first repetition unit. This fade-in of white noise ensured participants could clearly hear the initial countdown. The volume of the auditory stimuli was set to the loudest comfortable volume for each participant at the beginning of the experiment. We also confirmed that participants could clearly hear the tones.

### GVS

Following the standard GVS method (Lopez & Cullen, 2024), disposable electrodes (Kendall H124SG; Cardinal Health, Dublin, OH, USA; 30 mm × 24 mm) were attached to the skin over the left and right mastoid processes after alcohol disinfection. The assignment of anodes and cathodes to the left and right mastoid processes was balanced across participants. Electrical stimulation was generated by an electrical generator (SEN-8203, Nihon Kohden, Tokyo, Japan) and delivered to the electrodes via an electrical isolator (SS-104JMG, Nihon Kohden).

In all experiments, the perceptual threshold for each participant was assessed at the beginning (see the *Experimental Paradigm* section below), and the maximum current was set to 90% of that threshold. During trials of the GVS condition in the experimental session, current was applied in synchrony with the beat timing. The time profile of the current is described by the following equation (1), where the onset of each beat is set as time *t* = 0:

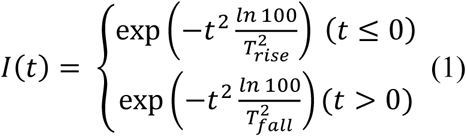

Equation (1) represents an asymmetric, monophasic Gaussian waveform, where *T_rise_* and *T_fall_* designate the time widths of the current rise and fall, respectively. *I* corresponds to the ratio relative to each participant’s 90% perceptual threshold. It is designed to reach 1 at the onset of the beat and to be 0.01 (i.e., 0.9% of the threshold) at both *T_rise_* seconds before and *T_fall_* seconds after that onset. According to studies showing that vestibular nerves exhibit a higher firing rate at the onset of GVS (Forbes, Kwan, Mitchell, Blouin, & Cullen, 2023), and when exposed to higher angular acceleration (Goldberg & Fernandez, 1971), which is equivalent to a larger time derivative of GVS when it imitates angular velocity (Gensberger et al., 2016), we manipulated the slope of the current rise. For experiment 1, we generated a waveform with a current rise time of 20 ms and a fall time of 100 ms by setting *T_rise_* and *T_fall_* to 0.02 and 0.1, respectively (Figure **1a**). For experiment 2, we flipped the waveform around its peak (*T_rise_* = 0.1, *T_fall_* = 0.02, Figure **1a**). By swapping the time widths of the current rise and fall between experiments, the total duration of the current application for the experimental session was controlled across experiments. At the same peak current, these two waveforms would produce identical peak magnitudes and spatial distributions of the electric field and current density in the skin beneath the electrodes, with temporally reversed profiles (Truong et al., 2024; Wang et al., 2024). By contrast, simulations based on the transfer functions reported by Allred et al. (2024) predicted distinct firing-rate dynamics in response to the two waveforms (Supplementary Figure S2). Therefore, this waveform design allowed us to examine the effect of the GVS parameter related to vestibular afferent firing on SMS performance.

**Figure 1.**
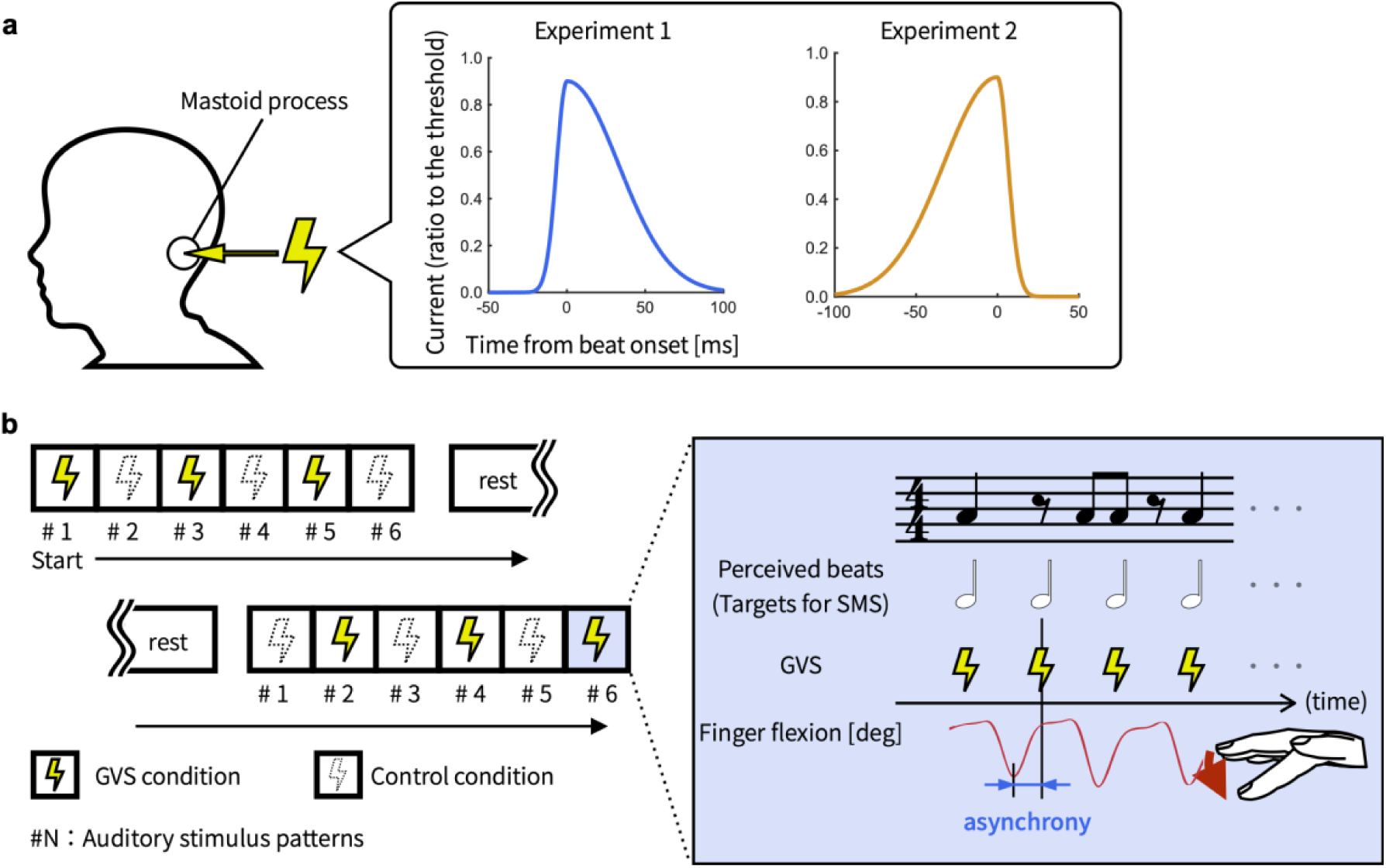
Design of the experiments. (**a**) Comparison of the current waveform between experiments 1 and 2. (**b**) Design of the experimental session, stimulus presentation, and behavioral responses. The left half depicts the block design of the experimental session. The blue-shaded area on the right illustrates the time course of stimulus presentation [auditory rhythms, perceived beats, and galvanic vestibular stimulation (GVS) shots]) and finger flexion.

### SMS Task

In the experiments, participants performed an SMS task; they flexed the index finger in synchrony with the beats in auditory rhythms. The practice session consisted of two phases: (1) finger flexion without an auditory stimulus and (2) flexion with an auditory stimulus. The first phase began with tapping on the surface of a height-adjustable platform. Its height was adjusted so that the fingertip touched the surface when the metacarpophalangeal (MCP) joint was flexed approximately 30 degrees. Moving from the baseline position with the index finger extended horizontally, participants practiced the proper tapping motion (quickly flexing the index finger at the MCP joint and then quickly extending it back to the baseline) to learn the appropriate movement and flexion angle. Next, after removing the platform, participants practiced finger flexion in the air, in the same manner as during tapping on the platform. The aims of this first phase were (1) to ensure that participants could consistently flex their index finger to 30 degrees and (2) ensure that they recognized reaching the end point of flexion as an event to be synchronized with the beat. In typical SMS with table tapping, the tactile input from the surface contact of the fingertip is used as sensory feedback for synchronization (Keller, Dalla Bella, & Koch, 2010; Keller, Ishihara, & Prinz, 2011). However, the present SMS task did not elicit such tactile input. Therefore, this phase was essential for participants to learn to use the somatosensory input at the end point of index finger flexion as sensory feedback for synchronization. In the second phase, participants listened to syncopation patterns and practiced index finger flexion to the beats. To determine the optimal syncopation intensity for each participant, when a participant achieved complete synchronization in a trial, the level was raised by 1 in the next trial, whereas it was reduced when a participant failed. Participants who failed in a trial with level 1 syncopation were excluded from the following analyses. Each participant completed at least five practice trials, with the option of additional trials if desired.

A standardized procedure was employed for the second phase of the practice and experimental sessions. The instructions given to participants were as follows: “Sit with your back straight, open your right hand so that the index finger and the sensor attached to it do not touch other parts of your hand, extend the index finger horizontally, wear the blindfold, do not move any body part other than your right index finger during the task, do not count the number of beats, do not memorize the rhythm by substituting words or phrases, do not recall any musical pieces, and concentrate on the task without thinking about anything else.” When the auditory stimulus began playing in each trial, participants immediately started synchronization by flexing the index finger to the countdown tones. Following those tones, a two-bar syncopation pattern played in a continuous loop, and participants maintained synchronization to the beat until the stimulus ended. Four and eight repetitions of a two-bar pattern were presented in each trial for the practice and experimental session, respectively. A single trial lasted approximately 45 s in the experimental session. Participants wore blindfolds while engaged in the SMS task.

### Experimental Paradigm

After participants arrived at the laboratory and completed the informed consent form and handedness questionnaire, preparations for the experiment began. The participant’s index finger on the right hand was secured in an extended position by applying tape from the MCP joint to the nail. Two wireless inertial sensors (WaveTrack Inertial System; Cometa Systems, Milan, Italy) were attached in a straight line on the dorsal surface of the right hand—one over the second metacarpal bone and one over the second proximal phalanx—so that the MCP joint was centered between these two sensors. The electrodes for GVS were attached to the skin over the mastoid processes (see the *GVS* section above). In addition, two wireless electromyography sensors (PicoEMG; Cometa Systems) were attached to the skin adjacent to these electrodes to monitor the applied electrical stimulation in real time.

Before the practice session, we conducted the procedure for estimating the perceptual threshold for GVS. First, we presented a train of waveforms identical to those used in each experiment, with a peak current of 100 µA, composed of three waves with an interval of 2 s. Participants were asked to report whether and how they experienced perceptual events during GVS (e.g., tingling or itching sensation on the skin beneath the electrodes, visual sway, or body tilt). We judged that perception had occurred only when the reports were consistent with the characteristics of the presented stimulus (e.g., the periodicity, intensity, location, or number of perceptual events). When the reports of perception did not correspond to the applied stimulus, we repeated application by increasing the amplitude of the current waveform in a fixed step size until correct perception was reported. Conversely, when the stimulus was correctly perceived, we reduced the amplitude in the same step size until no perception was reported. In other words, GVS was attenuated when perceived, whereas it was amplified when not perceived. The size of the current step started at 100 µA, and was halved when the second and fourth switches in step direction occurred, eventually being set to 25 µA. GVS with a peak current of 25 µA continued until the step direction switched six times. The individual perceptual threshold was defined as the mean amplitude at which those six switches occurred (Soranzo & Grassi, 2014; Wilkinson, Ko, Kilduff, McGlinchey, & Milberg, 2005). Note that the determined thresholds did not significantly differ between the experiments [a two-tailed one-sample t-test; *p* = 0.600, 95% confidence interval (CI) = –0.137, 0.233, *r* = 0.09].

After the perceptual threshold estimation, the practice and experimental sessions were conducted. Participants sat on a backless stool during the sessions. The experimental session consisted of two blocks of six trials, where GVS and control conditions alternated within each block (Figure **1b**). The conditions for the first trial in the first block were randomized across participants to ensure equal distribution. At the same time, the conditions assigned to the first trial in each block differed from one another. Six different syncopation patterns were assigned to each trial in the same order across both blocks. This design controlled the order effect for pattern presentation while assigning both conditions to each pattern. A break of ≥ 3 min was scheduled between blocks, and participants could take additional breaks at any time between trials on request.

### Data Recording

We controlled data recording and signal generation with an identical analog-to-digital (AD) converter (USB-6212 (BNC); National Instruments, Austin, TX, USA). This AD converter generated and output three signals: the GVS current waveform, auditory stimulus, and trigger signal for inertial data recording. These output signals were input to the electrical stimulus generator, headphones (MDR-7506; Sony, Tokyo, Japan), and an inertial data recording system (EMG and Motion Tools, version 7.4.6.0; Cometa Systems), respectively, and simultaneously input to the AD converter itself. These settings achieved precise temporal synchronization between stimulus presentation and data recording. We controlled the AD converter and data recording with the computational software MATLAB (R2023a, version 9.14.0.2286388; MathWorks, Natick, MA, USA) installed on a desktop PC. The software for recording inertial data was installed on another PC, where the inertial data were recorded. The acquisition of inertial data began immediately upon trigger input and was not affected by delays. The sampling frequency of the AD converter was 32,000 Hz, and that for inertial data was 284 Hz.

### Data Processing

Angular displacement data for index finger flexion were low-pass filtered at 6 Hz with a 4th-order Butterworth filter. We extracted the local peak of angular displacement closest to each beat with the MATLAB *findpeaks* function. The extracted peaks were defined as the onsets of movement synchronized to the beats. Before quantitative analyses, we confirmed that peak detection was correct by visual inspection. If the detection was incorrect, we adjusted the *findpeaks* parameters. For each trial, we included the onset of finger flexion to 64 beats within the syncopated rhythm in the analysis, whereas flexion to the countdowns was excluded. We calculated the temporal difference between the onset of each auditory beat and that of the corresponding finger flexion, which was defined as asynchrony (error of synchronization). Then, we calculated the absolute value and standard deviation (SD) of asynchrony for each trial as indices of accuracy and precision, respectively; SMS with higher accuracy and stability would result in a smaller absolute value and a smaller SD.

### Statistical Analyses

To eliminate the influence of differences in rhythm patterns across trials and between individuals, we compared the indices between the trials of the GVS condition and those of the control condition sharing the same syncopation pattern. We calculated Δ asynchrony and Δ |asynchrony| as indices of asynchrony and its absolute value, respectively, and assessed whether they were modulated in GVS trials compared with control trials. We subtracted the values of control trials from those of the corresponding GVS trials, and averaged the six resulting values from six pairs of trials within each individual. A two-tailed one-sample t-test examined whether the mean of Δ asynchrony or Δ |asynchrony| differed significantly from 0 across participants. Regarding the SD of asynchrony, we calculated the SD ratio within individuals as follows: we divided the SD values of GVS trials by those of the corresponding control trials, and averaged the six quotients for the six rhythm patterns within each participant. After applying natural logarithmic transformation, we used a two-tailed one-sample t-test to confirm whether the mean of the transformed SD ratio differed significantly from 0 across participants. The significance level for the t-test was set to 0.05. Intraindividual mean values identified as outliers by MATLAB’s *isoutlier* function were replaced with the mean value for participants without such outliers. Shapiro–Wilk tests confirmed the normality of the data distribution. Additionally, we reported the effect size (*r*; equivalent to η in one-way analysis of variance) for each t-test (Cohen, 1988; Hatch & Lazaraton, 1991).

## Results

### Experiment 1: GVS With a Steeper Current Increase Stabilized SMS Timing

In experiment 1, we applied a GVS waveform with a 20 ms rise time and a 100 ms fall time. The descriptive statistics are summarized in Table 1. Asynchrony SD was significantly reduced in the GVS condition compared with the control condition (*p* = 0.026, 95% CI = –8.37, –0.61, *r* = 0.48, Figure **2b**). The other two indices did not exhibit significant differences between conditions (asynchrony, *p* = 0.172, 95% CI = –1.00, 5.25, *r* = 0.31, Figure **2a**; absolute value of asynchrony, *p* = 0.063, 95% CI = –5.30, 0.15, *r* = 0.41, Figure **2c**). A previous study using a similar stimulation timeline (six 30 s GVSs at 1 min intervals) reported significant differences in behavioral outcomes between measurements before the first intervention and after the last one (Inukai, Miyaguchi, Saito, Otsuru, & Onishi, 2020). Given that GVS in the present experiment also activated the task-related neural substrates by the same mechanism, the long-term effect of GVS on performance would have remained throughout the control condition trials immediately following the GVS condition, resulting in no differences between conditions. Therefore, the results in experiment 1 indicate that GVS simultaneous with musical beats had an immediate timing-stabilizing effect on SMS to the beats rather than a long-term effect.

**Figure 2.**
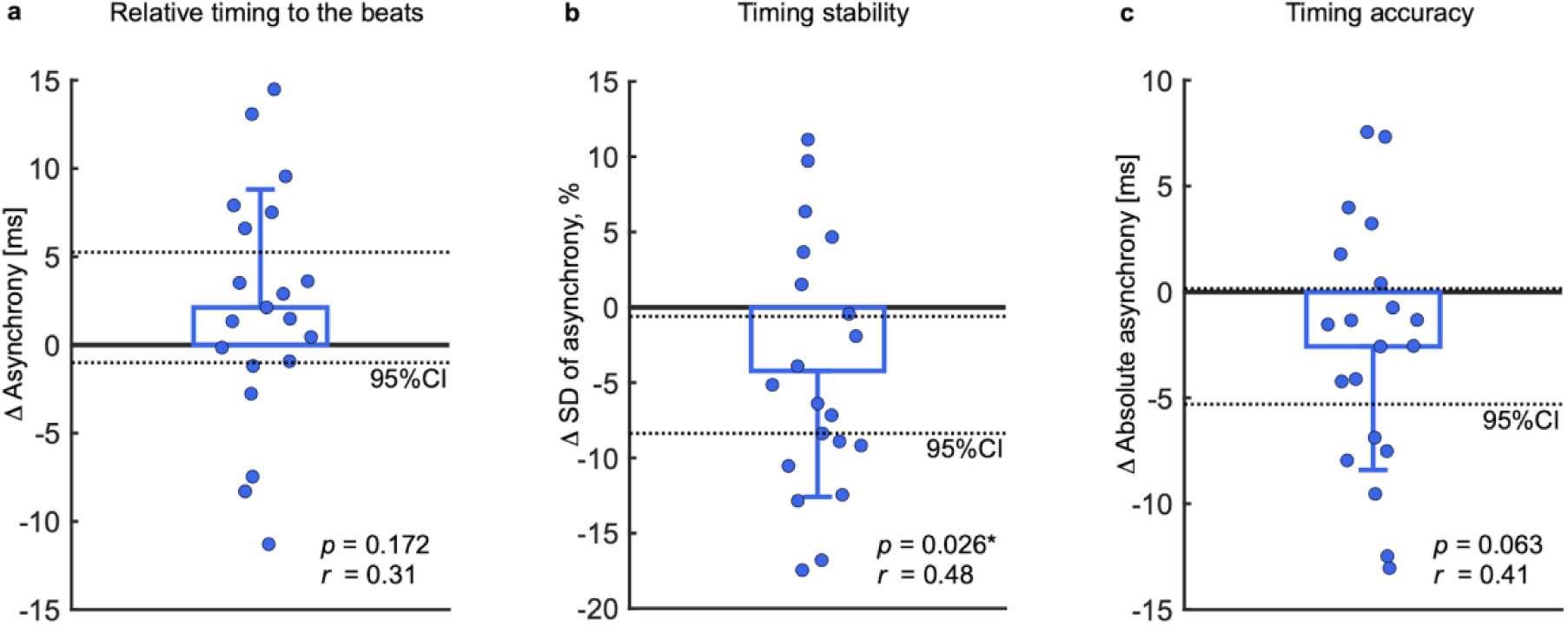
Sensorimotor synchronization (SMS) performance in trials with galvanic vestibular stimulation (GVS) compared with trials without GVS in experiment 1. (a) Asynchrony in GVS condition trials relative to control condition trials (Δ asynchrony). (b) Standard deviation (SD) of asynchrony in GVS trials relative to the SD in control trials (SD ratio). (c) Absolute value of asynchrony in GVS trials relative to control trials (Δ |asynchrony|). Each dot depicts a single participant (mean value within individuals). Bars and error bars illustrate the mean and standard deviation of the datapoints, respectively. The dotted lines indicate the 95% confidence intervals for the t-test. p* < 0.05.

**Table 1.**
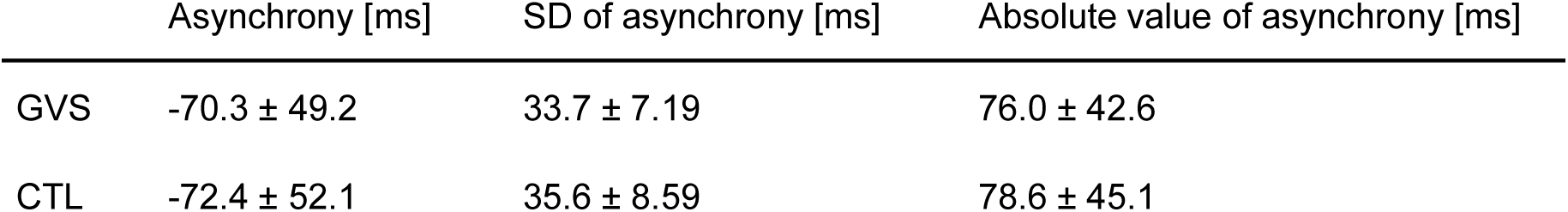
Descriptive statistics of the data from experiment 1. The values represent the grand average (± standard error of the mean) for each condition. GVS, galvanic vestibular stimulation condition; CTL, control condition.

|  | Asynchrony [ms] | SD of asynchrony [ms] | Absolute value of asynchrony [ms] |
| --- | --- | --- | --- |
| GVS | -70.3 ± 49.2 | 33.7 ± 7.19 | 76.0 ± 42.6 |
| CTL | -72.4 ± 52.1 | 35.6 ± 8.59 | 78.6 ± 45.1 |

### Experiment 2: GVS With a Gradual Current Increase Had No Influence on SMS Timing

The results of Experiment 1 suggested that vestibular input synchronized with musical beats enhanced the precision of SMS to those beats. However, we considered this phenomenon alone to be insufficient to support the hypothesis that the vestibular system plays a functional role in the execution of predictive movements to musical rhythms. Previous studies reported that SMS was more stable when multimodal rhythmic cues were presented, such as auditory and visual cues, compared with auditory cues alone (Elliott, Wing, & Welchman, 2010; Roy, Dalla Bella, & Lagarde, 2017; Wing, Doumas, & Welchman, 2010). Therefore, we could not rule out the possibility that, similar to the findings for visual and tactile cues in those previous studies, GVS served only as an additional sensory modality providing rhythmic cues, resulting in SMS stabilization. We addressed this issue in experiment 2 (the control experiment) using a flipped waveform with a rise time of 100 ms and a fall time of 20 ms. The descriptive statistics are summarized in Table 2. There was no significant difference in the SD of asynchrony between the GVS and control conditions (*n* = 20; *p* = 0.482, 95% CI = –4.00, 8.70, *r* = 0.16, Figure **3b**), and no significant changes were observed in other indices (asynchrony, *p* = 0.914, 95% CI = –2.85, 3.16, *r* = 0.03, Figure **3a**; absolute value of asynchrony, *p* = 0.901, 95% CI = –4.60, 5.19, *r* = 0.03, Figure **3c**). These results indicate that merely applying GVS in synchrony with the beats does not positively affect SMS, instead demonstrating that the effect of GVS on SMS could depend on the GVS parameters. Although the total duration and amplitude of the applied current did not differ from experiment 1, the effect of GVS on SMS performance disappeared in experiment 2. The data suggest that the temporal structure of the current waveform likely determined the effect. Experiment 2 also corroborated that the timing-stabilizing effect of GVS in experiment 1 was not attributable to the addition of a sensory modality providing timing cues, as in previously reported multimodal SMS (Elliott et al., 2010; Roy et al., 2017; Wing et al., 2010).

**Figure 3.**
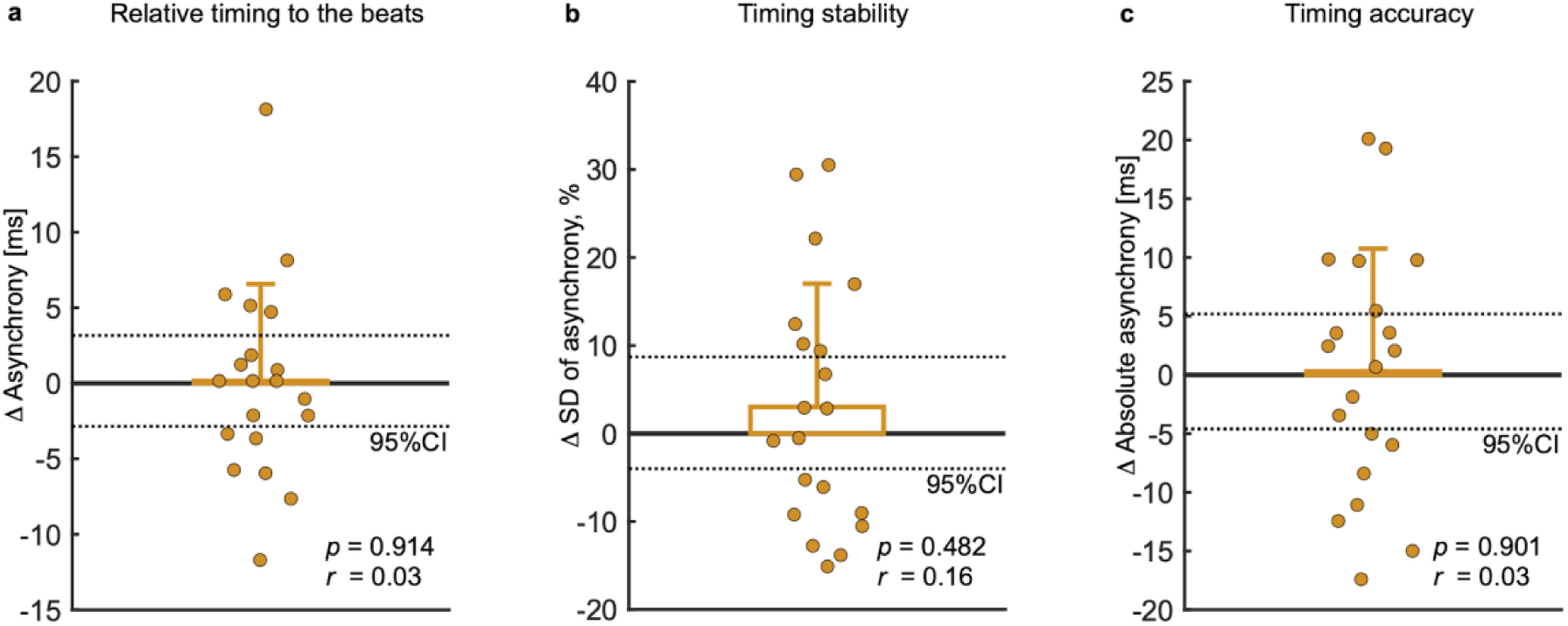
Sensorimotor synchronization (SMS) performance in trials with galvanic vestibular stimulation (GVS) compared with trials without GVS in experiment 2. (**a**) Asynchrony in GVS condition trials relative to control condition trials (Δ asynchrony). (**b**) Standard deviation (SD) of asynchrony in GVS trials relative to the SD in control trials (SD ratio). (**c**) Absolute value of asynchrony in GVS trials relative to control trials (Δ |asynchrony|). Each dot depicts a single participant (mean value within individuals). Bars and error bars illustrate the mean and standard deviation of the datapoints, respectively. The dotted lines indicate the 95% CIs used for t-test.

**Table 2.** Descriptive statistics of the data from experiment 2. The values represent the grand average (± standard error of the mean) for each condition. GVS, galvanic vestibular stimulation condition; CTL, control condition.

|  | Asynchrony [ms] | SD of asynchrony [ms] | Absolute value of asynchrony [ms] |
| --- | --- | --- | --- |
| GVS | -72.5 ± 43.1 | 40.5 ± 10.75 | 75.4 ± 37.2 |
| CTL | -72.6 ± 41.8 | 39.3 ± 9.04 | 75.1 ± 34.7 |

## Discussion

The present study demonstrated for the first time, using behavioral data from healthy adults, that the vestibular system is involved in SMS, a primitive and complex behavior requiring sensory pattern detection and predictive motor control. There has been a debate for decades on the significance of the behavior (or habit) of moving the head or body in synchrony with musical rhythms (Todd & Lee, 2015). However, direct investigation of this issue has been limited owing to the difficulty of measuring neural activity within healthy humans during head movements. To address this issue, this study employed a non-invasive method to stimulate the vestibular system and examined the influence of vestibular input generated by head movements on predictive motor execution anticipatory of periodic musical beats. Our data provided causal evidence that vestibular input contributes immediately to SMS timing stabilization, which suggests that vestibular system involvement underlies the benefits of head nodding and body sway for SMS performance.

In experiment 1, a GVS with a current rise time of 20 ms and a fall time of 100 ms yielded an immediate improvement in SMS timing stability. The most straightforward interpretation of this result is that the congruence in periodicity between the beats and the vestibular inputs enhanced beat perception, leading to better SMS. It has already been confirmed that perception for an ambiguous meter structure is biased toward the periodicity of the vestibular input experienced while listening to such a structure (Trainor et al., 2009). Our GVS with the same periodicity as the beats may also have facilitated beat perception. In particular, beats in syncopation were difficult to perceive (Chapin et al., 2010; Tal et al., 2017), and SMS with syncopation is difficult and inferior compared with that with an isochronous metronome (Fitch & Rosenfeld, 2007; Matthews, Witek, Thibodeau, Vuust, & Penhune, 2022; Patel, Iversen, Chen, & Repp, 2005; Witek et al., 2014). Therefore, such enhanced beat perception may have improved the precision of SMS with syncopated rhythms.

However, there remains a discrepancy in that the effect of GVS in experiment 1 was instantaneous, and GVS had little long-term effect (thus not affecting the subsequent trials), whereas the effect of GVS on beat perception was achieved through training in previous studies (Tichko et al., 2021; Trainor et al., 2009). This prompt effect could be explained to some extent by the mechanism whereby multimodal timing signals improve SMS stability (Elliott et al., 2010; Roy et al., 2017; Wing et al., 2010). For example, Roy et al. (2017) reported that SMS stability improved when the time difference between tactile and auditory events was congruent with the difference in perceptual latencies (i.e., when the two sensory events were consciously perceived as simultaneous). Importantly, in the present study, GVS was applied at subthreshold intensity, inducing no perceptual events. Thus, it is unclear whether the vestibular input in this study functioned similarly to the tactile input in previous studies because the GVS was not consciously perceived in our study. Accordingly, it was necessary to further verify whether the improvement in SMS in experiment 1 was solely due to the co-existence of auditory and vestibular inputs. To address this, we altered the current waveform in the subsequent experiment.

In experiment 2, the temporal profile of the GVS waveform was flipped back and forth in time, allowing us to control the total amount of current applied across experiments. Supposing that the congruence of periodicity between the beats and GVS improved SMS stability in experiment 1, we expected to obtain a similar result with this flipped waveform. However, the waveform in experiment 2 had no significant effect on SMS, suggesting specificity of SMS modulation to GVS parameters other than the timing of application or the total applied current. It was also confirmed that the SMS stabilization in experiment 1 was not caused solely by the addition of sensory modalities providing timing cues. To explain the inconsistency of the influence of GVS on performance between experiments, we focused on the differences in waveform between experiments: how steeply the current rose, and when the current changed relative to the beats.

We interpreted the results from the perspective of the steepness of the current onset. In other words, we speculated that the size of the time derivative of the current for onsets might be the determinant of SMS stabilization. This idea is supported by the firing dynamics of vestibular afferent fibers, which exhibit a higher firing rate during exposure to higher angular acceleration (Goldberg & Fernandez, 1971). Because angular acceleration is the time derivative of angular velocity, and the GVS current corresponds to the angular velocity of head rotation (Gensberger et al., 2016), the steepness of the current rise could impact the firing rate similarly to the angular acceleration of head rotation. Accordingly, the waveform used in experiment 1 might have had a sufficient slope to induce afferent fiber firing affecting performance, unlike the waveform in experiment 2. In addition, it is reasonable that the effect of GVS depends on waveform onset because the dynamics of the firing rates of vestibular afferent fibers are asymmetrical between the onset and offset of current application (Forbes et al., 2023). Therefore, it is reasonable to ascribe the inconsistency in the GVS effect between the experiments to the steepness of the current rise.

Given the temporal relationship between vestibular activation and beat onset, we can further discuss a possible underlying mechanism: the interaction with the thalamus, one of the primary relays for vestibular information (Lee, Park, & Yeo, 2025; Lopez & Blanke, 2011; Wijesinghe, Protti, & Camp, 2015). The central thalamus was recently revealed to be involved in the prediction of rhythmic sensory inputs by encoding their timing (Matsuyama & Tanaka, 2021). That research demonstrated that when specific neurons in the central thalamus exhibit increased activity during a 100 ms time window before rhythmic events, the latency of movements toward those events is shortened. Moreover, the firing dynamics of those thalamic neurons were similar to the typical temporal profile of asynchrony in SMS (Takeya, Kameda, Patel, & Tanaka, 2017). Given that peripheral vestibular afferent signals reach the thalamus in approximately 2.5–5 ms (Deecke, Schwarz, & Fredrickson, 1974; Liedgren, Milne, Schwarz, & Tomlinson, 1976), it is conceivable that the GVS in experiment 1 might have activated thalamic neurons before beat onset, promoting predictive encoding for beat timing and improving the stability of SMS.

Not only the thalamus, but also other cortical and subcortical regions, might have received vestibular projections triggered by GVS (Nakul, Bartolomei, & Lopez, 2021; Stiles & Smith, 2015). Given that vestibular stimulation causes evoked potentials in various cortical regions, including frontal, parietal, temporal and occipital areas, within a short latency of 10 or 20 ms (Nakul et al., 2021), GVS concurrent with the beats could have led to entrained neural activity within those areas. In fact, Rosso, Leman, and Moumdjian (2021) revealed that higher accuracy and higher stability were accompanied by more stable neural oscillation at the beat frequency recorded from the post-frontocentral area. In addition, the spectral power of neural oscillation at the beat frequency correlated with SMS performance and the accuracy of beat prediction (Nozaradan, Peretz, & Keller, 2016). These findings highlight the tight connection between SMS and rhythmic neural activity at the beat frequency. Accordingly, GVS at beat onset might have enhanced neural activity at the same frequency as the beats in both experiments. However, SMS was promoted in the GVS condition only in experiment 1. Hence, the entrainment of cortical activity might not be the primary factor underlying the stabilization effect of GVS.

The GVS procedure in this study was novel, using an asymmetric, monophasic Gaussian-shaped current waveform and flipping the waveform across experiments. The typical waveforms of GVS current include pulse, sine, and noise waves (Dlugaiczyk, Gensberger, & Straka, 2019). Pulse waves are often applied in the form of trains at frequencies ranging from tens to hundreds of Hz, and each pulse lasts for microseconds (Pfanzelt et al., 2008; Phillips et al., 2015). Sinusoidal GVS is typically applied continuously at 0.01 to tens of Hz over hundreds of seconds, and each sine wave is biphasic (Cohen et al., 2011; Kwan, Forbes, Mitchell, Blouin, & Cullen, 2019; Macefield & James, 2016). Noise GVS is composed of zero-mean alternating noise (usually white noise) waveforms with durations of 5 s to 30 min (McLaren, Smith, Taylor, Niazi, & Taylor, 2023). Even though a few studies used symmetrical, biphasic Gaussian-shaped waveforms 500 ms in width (Barnett-Cowan & Harris, 2009; Trainor et al., 2009), our GVS method was challenging in terms of the shape of the waveform, the instantaneous and intermittent application, and the time-lock with other sensory events. Nevertheless, our GVS successfully modulated behavioral performance requiring predictive motor control and auditory perceptual processing. Furthermore, the asymmetry of the waveform played an important role in interpreting the mechanisms underlying the results. Altogether, our results emphasize the significance of designing GVS waveforms according to the aims of the study and extend the framework of GVS research.

The SMS stabilization observed in experiment 1 was consistent with our previous study, where SMS was stabilized when moving the head to the beats during the same task as in the present study (Yamazaki & Ushiyama, 2024). Owing to the abundance of neck muscle spindles (Banks, 2006; Kissane, Charles, Banks, & Bates, 2023), the possibility that the proprioceptive feedback from neck flexion was the primary factor improving SMS in the finger cannot be ruled out (Bravi, Cohen, Martinelli, Gottard, & Minciacchi, 2017). Therefore, we did not let the participants move their heads and used unperceivable stimulation in the present study, whose results allowed us to rule out any contribution of proprioceptive feedback. Taken together, these two studies suggest that SMS stabilization related to simultaneous head movements could be caused by the vestibular input accompanying the head movements. In addition, even a short-duration, low-intensity (subthreshold) GVS could potentially elicit neural activity homogeneous with that arising during head movement. This possibility points not only to the safety of our novel GVS technique but also to its potential as a non-invasive intervention simulating vestibular activities during head movements, highlighting the utility of the technique for fundamental research and clinical applications.

Although the present study provides the first causal evidence for direct involvement of the vestibular system in timing behavior anticipatory of musical rhythms, it has several limitations. The primary limitation is that the dynamics of afferent fiber firing in the semicircular canals and otolith organs triggered by GVS is generally not the same as that during natural head movements (Lopez & Cullen, 2024). However, owing to the brief application (120 ms) around each beat onset, the vestibular signals elicited by GVS might have been more similar to those accompanied by head movements to music than is seen with the typical methods discussed above. Future research should investigate the effects of other GVS parameters, such as electrode placement (Aoyama, Iizuka, Ando, & Maeda, 2015), rather than the slope of the current waveform.

Various parameters or components of the SMS task should also be taken into account, such as the tempo of the rhythm. The interval between perceived beats in this study was 600 ms (i.e., 1.67 Hz), which is close to the frequency of vertical acceleration of the head in daily life (approximately 2 Hz; MacDougall & Moore, 2005). Therefore, there is room to investigate whether the beneficial effect of vestibular input on motor performance might be specific to the frequency that is actually experienced (or to a frequency on which a participant has been well trained). The present results might be specific to syncopation, which has a greater requirement for beat prediction than a metronome (Vuust, Dietz, Witek, & Kringelbach, 2018; Vuust et al., 2022; Zalta, Large, Schön, & Morillon, 2024). In addition to these task characteristics, participants’ musical background (e.g., the onset, years, or content of music or dance training) could have influenced the results (Baer, Thibodeau, Gralnick, Li, & Penhune, 2013; Bailey & Penhune, 2010; Cirelli, Spinelli, Nozaradan, & Trainor, 2016; Giacosa, Karpati, Foster, Penhune, & Hyde, 2016; Jin et al., 2019; Karpati, Giacosa, Foster, Penhune, & Hyde, 2016; Li et al., 2015; Matthews, Thibodeau, Gunther, & Penhune, 2016; Miura, Fujii, Okano, Kudo, & Nakazawa, 2016; Miura, Kudo, Ohtsuki, & Kanehisa, 2011; Roman, Washburn, Large, Chafe, & Fujioka, 2019). Even though these factors were out of the scope of the present study, their involvement should be clarified in future studies.

In conclusion, this study provides the first causal evidence that the vestibular system has a functional role in movements synchronized to musical rhythms. This finding offers a new perspective on the behaviors of head bobbing and body swaying to musical rhythms observed across cultures and animal species (Fitch, 2013; Palmer et al., 2019; Schachner et al., 2009; Swarbrick et al., 2019; Van Dyck et al., 2013). The results suggested that the size of the GVS time derivative is one of the determinants of the vestibular contribution to predictive motor execution anticipatory of upcoming auditory events. This finding deepens our comprehension of the vestibular system and the circuits integrating the information that it provides. In addition, considering that the results were obtained using our novel GVS method with subthreshold and short-duration waveforms, the present study emphasizes the importance of GVS parameters other than duration and intensity.

## Acknowledgements

The study participants were recruited via a website designed for the recruitment of experimental participants (https://www.jikken-baito.com). We thank Ms. Tomomi Hamaoka for her secretarial assistance and all other members of our laboratory for their insightful comments on the work. We also thank Michael Irvine, from Edanz (https://www.edanz.com/ac) for editing a draft of this manuscript.

## Author contributions

J.U. supervised the research, and reviewed and edited the manuscripts. R.Y. curated and analyzed the data, investigated articles, developed the code, visualized the data, and wrote the original draft. All authors conceptualized the research, acquired funding, established the methodology, and managed the project.

## Data availability

The Ethics Committee of our institute permitted the present research under the following condition: “The data retention period is set at ten years from the publication of the original research paper in an academic journal. After this period, all electronic and paper-based data must be destroyed.” Therefore, the data obtained in the present experiments are not publicly available, although they are available upon request and with the authors’ permission.

---

**Supplementary Figure S1.**
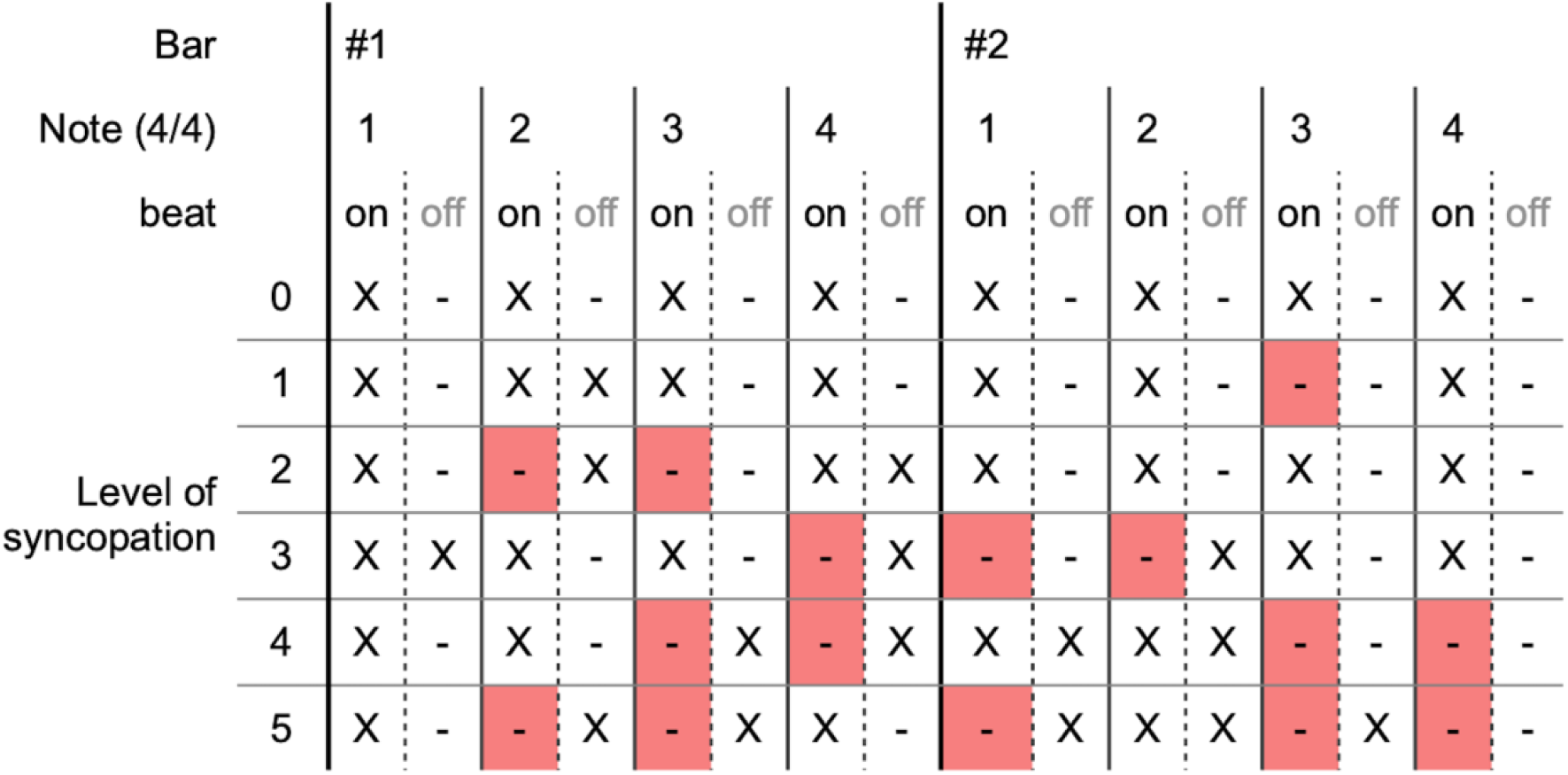
Examples of syncopated rhythm patterns. “X” denotes the placement of each tone, whereas “-” denotes the absence of tones. “Level of syncopation” corresponds to the number of on-beat (namely, tones in an isochronous metronome) to off-beat replacements; the pattern of level 0 is identical to metronome sounds. Red-shaded cells indicate the places where on-beat tones were removed.

**Supplementary Figure S2.**
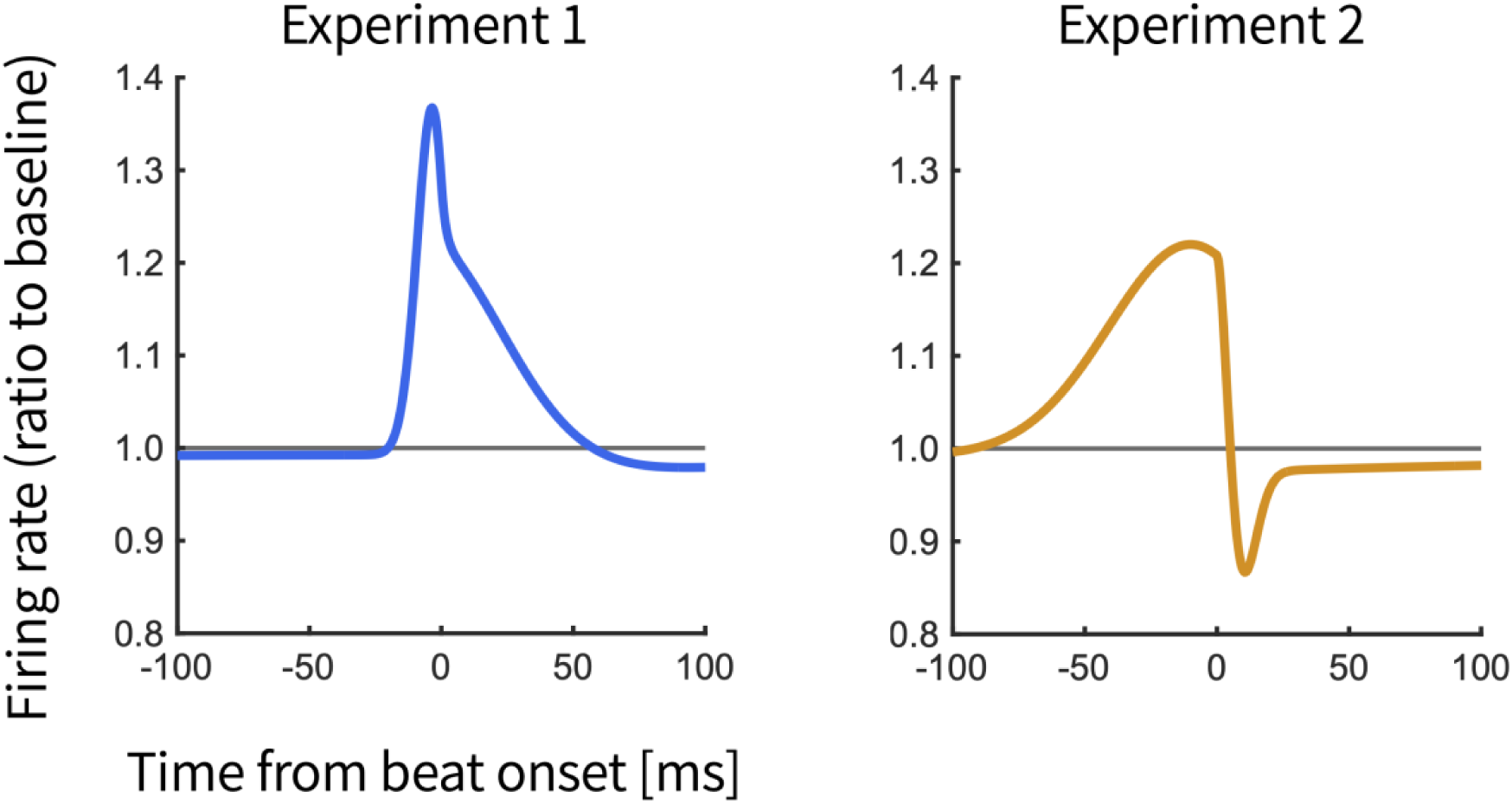
Simulation of vestibular afferent firing induced by GVS. The left and right panels depict the predicted temporal dynamics of the firing rate of vestibular afferent nerves for semicircular canals on the cathodal side in response to the GVS current application in experiments 1 and 2, respectively. The x-axis represents time relative to beat onsets in syncopated rhythms. The firing rate of baseline is normalized to 1 (represented by the gray horizontal lines). The simulation was based on the transfer function for irregular afferent nerves of semicircular canals reported by Allred et al. (2024).

## References

Allred, A. R., Austin, C. R., Klausing, L., Boggess, N., & Clark, T. K. (2024). Human perception of self-motion and orientation during galvanic vestibular stimulation and physical motion. PLOS Computational Biology, 20(11), e1012601. 10.1371/journal.pcbi.1012601

Aoyama, K., Iizuka, H., Ando, H., & Maeda, T. (2015). Four-pole galvanic vestibular stimulation causes body sway about three axes. Scientific Reports, 5, 10168. 10.1038/srep10168

Baer, L. H., Thibodeau, J. L. N., Gralnick, T. M., Li, K. Z. H., & Penhune, V. B. (2013). The role of musical training in emergent and event-based timing. Frontiers in Human Neuroscience, 7, 191. 10.3389/fnhum.2013.00191

Bailey, J. A., & Penhune, V. B. (2010). Rhythm synchronization performance and auditory working memory in early- and late-trained musicians. Experimental Brain Research, 204, 91–101. 10.1007/s00221-010-2299-y

Banks, R. W. (2006). An allometric analysis of the number of muscle spindles in mammalian skeletal muscles. Journal of Anatomy, 208, 753–768. 10.1111/j.1469-7580.2006.00558.x

Barnett-Cowan, M., & Harris, L. R. (2009). Perceived timing of vestibular stimulation relative to touch, light and sound. Experimental Brain Research, 198, 221–231. 10.1007/s00221-009-1779-4

Bravi, R., Cohen, E. J., Martinelli, A., Gottard, A., & Minciacchi, D. (2017). When non-dominant is better than dominant: Kinesiotape modulates asymmetries in timed performance during a synchronization-continuation task. Frontiers in Integrative Neuroscience, 11, 21. 10.3389/fnint.2017.00021

Burger, B., Thompson, M. R., Luck, G., Saarikallio, S., & Toiviainen, P. (2013). Influences of rhythm- and timbre-related musical features on characteristics of music-induced movement. Frontiers in Psychology, 4, 183. 10.3389/fpsyg.2013.00183

Chapin, H. L., Zanto, T., Jantzen, K. J., Kelso, S. J. A., Steinberg, F., & Large, E. W. (2010). Neural responses to complex auditory rhythms: The role of attending. Frontiers in Psychology, 1, 224. 10.3389/fpsyg.2010.00224

Cirelli, L. K., Spinelli, C., Nozaradan, S., & Trainor, L. J. (2016). Measuring neural entrainment to beat and meter in infants: Effects of music background. Frontiers in Neuroscience, 10, 229. 10.3389/fnins.2016.00229

Cohen, B., Martinelli, G. P., Ogorodnikov, D., Xiang, Y., Raphan, T., Holstein, G. R., & Yakushin, S. B. (2011). Sinusoidal galvanic vestibular stimulation (sGVS) induces a vasovagal response in the rat. Experimental Brain Research, 210, 45–55. 10.1007/s00221-011-2604-4

Cohen, J. (1988). Statistical power analysis for the behavioral sciences (2nd ed.). Lawrence Erlbaum Associates. 10.4324/9780203771587

Deecke, L., Schwarz, D. W. F., & Fredrickson, J. M. (1974). Nucleus ventroposterior inferior (VPI) as the vestibular thalamic relay in the rhesus monkey. I. Field potential investigation. Experimental Brain Research, 20, 88–100. 10.1007/bf00239019

Dlugaiczyk, J., Gensberger, K. D., & Straka, H. (2019). Galvanic vestibular stimulation: From basic concepts to clinical applications. Journal of Neurophysiology, 121, 2237–2255. 10.1152/jn.00035.2019

Elliott, M. T., Wing, A. M., & Welchman, A. E. (2010). Multisensory cues improve sensorimotor synchronisation. European Journal of Neuroscience, 31, 1828–1835. 10.1111/j.1460-9568.2010.07205.x

Faul, F., Erdfelder, E., Lang, A.-G., & Buchner, A. (2007). G*Power 3: A flexible statistical power analysis program for the social, behavioral, and biomedical sciences. Behavior Research Methods, 39, 175–191. 10.3758/bf03193146

Fitch, W. T. (2013). Rhythmic cognition in humans and animals: Distinguishing meter and pulse perception. Frontiers in Systems Neuroscience, 7, 68. 10.3389/fnsys.2013.00068

Fitch, W. T., & Rosenfeld, A. J. (2007). Perception and production of syncopated rhythms. Music Perception, 25, 43–58. 10.1525/mp.2007.25.1.43

Forbes, P. A., Kwan, A., Mitchell, D. E., Blouin, J.-S., & Cullen, K. E. (2023). The neural basis for biased behavioral responses evoked by galvanic vestibular stimulation in primates. Journal of Neuroscience, 43, 1905–1919. 10.1523/jneurosci.0987-22.2023

Gensberger, K. D., Kaufmann, A.-K., Dietrich, H., Branoner, F., Banchi, R., Chagnaud, B. P., & Straka, H. (2016). Galvanic vestibular stimulation: Cellular substrates and response patterns of neurons in the vestibulo-ocular network. Journal of Neuroscience, 36, 9097–9110. 10.1523/jneurosci.4239-15.2016

Giacosa, C., Karpati, F. J., Foster, N. E. V., Penhune, V. B., & Hyde, K. L. (2016). Dance and music training have different effects on white matter diffusivity in sensorimotor pathways. NeuroImage, 135, 273–286. 10.1016/j.NeuroImage.2016.04.048

Goldberg, J. M., & Fernandez, C. (1971). Physiology of peripheral neurons innervating semicircular canals of the squirrel monkey. I. Resting discharge and response to constant angular accelerations. Journal of Neurophysiology, 34, 635–660. 10.1152/jn.1971.34.4.635

Hatch, E. M., & Lazaraton, A. (1991). The research manual: Design and statistics for applied linguistics (2nd ed.). Newbury House Publishers.

Inukai, Y., Miyaguchi, S., Saito, M., Otsuru, N., & Onishi, H. (2020). Effects of different stimulation conditions on the stimulation effect of noisy galvanic vestibular stimulation. Frontiers in Human Neuroscience, 14, 581405. 10.3389/fnhum.2020.581405

Jin, X., Wang, B., Lv, Y., Lu, Y., Chen, J., & Zhou, C. (2019). Does dance training influence beat sensorimotor synchronization? Differences in finger-tapping sensorimotor synchronization between competitive ballroom dancers and nondancers. Experimental Brain Research, 237, 743–753. 10.1007/s00221-018-5410-4

Karpati, F. J., Giacosa, C., Foster, N. E. V., Penhune, V. B., & Hyde, K. L. (2016). Sensorimotor integration is enhanced in dancers and musicians. Experimental Brain Research, 234, 893–903. 10.1007/s00221-015-4524-1

Keller, P. E., Dalla Bella, S., & Koch, I. (2010). Auditory imagery shapes movement timing and kinematics: Evidence from a musical task. Journal of Experimental Psychology: Human Perception and Performance, 36, 508–513. 10.1037/a0017604

Keller, P. E., Ishihara, M., & Prinz, W. (2011). Effects of feedback from active and passive body parts on spatial and temporal parameters in sensorimotor synchronization. Cognitive Processing, 12, 127–133. 10.1007/s10339-010-0361-0

Kissane, R. W. P., Charles, J. P., Banks, R. W., & Bates, K. T. (2023). The association between muscle architecture and muscle spindle abundance. Scientific Reports, 13, 2830. 10.1038/s41598-023-30044-w

Kwan, A., Forbes, P. A., Mitchell, D. E., Blouin, J.-S., & Cullen, K. E. (2019). Neural substrates, dynamics and thresholds of galvanic vestibular stimulation in the behaving primate. Nature Communications, 10, 1904. 10.1038/s41467-019-09738-1

Large, E. W., Herrera, J. A., & Velasco, M. J. (2015). Neural networks for beat perception in musical rhythm. Frontiers in Systems Neuroscience, 9, 159. 10.3389/fnsys.2015.00159

Lee, S.-S., Park, S.-Y., & Yeo, S.-S. (2025). Different types of connections between the thalamus and vestibular nucleus in the human brain. Journal of Clinical Medicine, 14, 7551. 10.3390/jcm14217551

Li, G., He, H., Huang, M., Zhang, X., Lu, J., Lai, Y., Luo, C., & Yao, D. (2015). Identifying enhanced cortico-basal ganglia loops associated with prolonged dance training. Scientific Reports, 5, 10271. 10.1038/srep10271

Liedgren, S. R., Milne, A. C., Schwarz, D. W., & Tomlinson, R. D. (1976). Representation of vestibular afferents in somatosensory thalamic nuclei of the squirrel monkey (*Saimiri sciureus*). Journal of Neurophysiology, 39, 601–612. 10.1152/jn.1976.39.3.601

Lopez, C., & Blanke, O. (2011). The thalamocortical vestibular system in animals and humans. Brain Research Reviews, 67, 119–146. 10.1016/j.brainresrev.2010.12.002

Lopez, C., & Cullen, K. E. (2024). Electrical stimulation of the peripheral and central vestibular system. Current Opinion in Neurology, 37, 40–51. 10.1097/wco.0000000000001228

MacDougall, H. G., & Moore, S. T. (2005). Marching to the beat of the same drummer: The spontaneous tempo of human locomotion. Journal of Applied Physiology, 99, 1164–1173. 10.1152/japplphysiol.00138.2005

Macefield, V. G., & James, C. (2016). Superentrainment of muscle sympathetic nerve activity during sinusoidal galvanic vestibular stimulation. Journal of Neurophysiology, 116, 2689–2694. 10.1152/jn.00036.2016

Matsuyama, K., & Tanaka, M. (2021). Temporal prediction signals for periodic sensory events in the primate central thalamus. Journal of Neuroscience, 41, 1917–1927. 10.1523/jneurosci.2151-20.2021

Matthews, T. E., Thibodeau, J. N. L., Gunther, B. P., & Penhune, V. B. (2016). The impact of instrument-specific musical training on rhythm perception and production. Frontiers in Psychology, 7, 69. 10.3389/fpsyg.2016.00069

Matthews, T. E., Witek, M. A. G., Thibodeau, J. L. N., Vuust, P., & Penhune, V. B. (2022). Perceived motor synchrony with the beat is more strongly related to groove than measured synchrony. Music Perception, 39, 423–442. 10.1525/mp.2022.39.5.423

McLaren, R., Smith, P. F., Taylor, R. L., Niazi, I. K., & Taylor, D. (2023). Scoping out noisy galvanic vestibular stimulation: A review of the parameters used to improve postural control. Frontiers in Neuroscience, 17, 1156796. 10.3389/fnins.2023.1156796

Miura, A., Fujii, S., Okano, M., Kudo, K., & Nakazawa, K. (2016). Finger-to-beat coordination skill of non-dancers, street dancers, and the world champion of a street-dance competition. Frontiers in Psychology, 7, 542. 10.3389/fpsyg.2016.00542

Miura, A., Kudo, K., Ohtsuki, T., & Kanehisa, H. (2011). Coordination modes in sensorimotor synchronization of whole-body movement: A study of street dancers and non-dancers. Human Movement Science, 30, 1260–1271. 10.1016/j.humov.2010.08.006

Nakul, E., Bartolomei, F., & Lopez, C. (2021). Vestibular-evoked cerebral potentials. Frontiers in Neurology, 12, 674100. 10.3389/fneur.2021.674100

Nicholls, M. E. R., Thomas, N. A., Loetscher, T., & Grimshaw, G. M. (2013). The Flinders Handedness Survey (FLANDERS): A brief measure of skilled hand preference. Cortex, 49, 2914–2926. 10.1016/j.cortex.2013.02.002

Nozaradan, S., Peretz, I., & Keller, P. E. (2016). Individual differences in rhythmic cortical entrainment correlate with predictive behavior in sensorimotor synchronization. Scientific Reports, 6, 20612. 10.1038/srep20612

Okubo, M., Suzuki, H., & Nicholls, M. E. R. (2014). A Japanese version of the FLANDERS handedness questionnaire. The Japanese Journal of Psychology, 85, 474–481. 10.4992/jjpsy.85.13235

Palmer, C., Spidle, F., Koopmans, E., & Schubert, P. (2019). Ears, heads, and eyes: When singers synchronise. Quarterly Journal of Experimental Psychology, 72, 2272–2287. 10.1177/1747021819833968

Patel, A. D., Iversen, J. R., Chen, Y., & Repp, B. H. (2005). The influence of metricality and modality on synchronization with a beat. Experimental Brain Research, 163, 226–238. 10.1007/s00221-004-2159-8

Pfanzelt, S., Rössert, C., Rohregger, M., Glasauer, S., Moore, L. E., & Straka, H. (2008). Differential dynamic processing of afferent signals in frog tonic and phasic second-order vestibular neurons. Journal of Neuroscience, 28, 10349–10362. 10.1523/jneurosci.3368-08.2008

Phillips, J. O., Ling, L., Nie, K., Jameyson, E., Phillips, C. M., Nowack, A. L., Golub, J. S., & Rubinstein, J. T. (2015). Vestibular implantation and longitudinal electrical stimulation of the semicircular canal afferents in human subjects. Journal of Neurophysiology, 113, 3866–3892. 10.1152/jn.00171.2013

Phillips-Silver, J., & Trainor, L. J. (2005). Feeling the beat: Movement influences infant rhythm perception. Science, 308, 1430. 10.1126/science.1110922

Phillips-Silver, J., & Trainor, L. J. (2007). Hearing what the body feels: Auditory encoding of rhythmic movement. Cognition, 105, 533–546. 10.1016/j.cognition.2006.11.006

Phillips-Silver, J., & Trainor, L. J. (2008). Vestibular influence on auditory metrical interpretation. Brain and Cognition, 67, 94–102. 10.1016/j.bandc.2007.11.007

Roman, I. R., Washburn, A., Large, E. W., Chafe, C., & Fujioka, T. (2019). Delayed feedback embedded in perception-action coordination cycles results in anticipation behavior during synchronized rhythmic action: A dynamical systems approach. PLOS Computational Biology, 15, e1007371. 10.1371/journal.pcbi.1007371

Rosso, M., Leman, M., & Moumdjian, L. (2021). Neural entrainment meets behavior: The stability index as a neural outcome measure of auditory-motor coupling. Frontiers in Human Neuroscience, 15, 668918. 10.3389/fnhum.2021.668918

Roy, C., Dalla Bella, S., & Lagarde, J. (2017). To bridge or not to bridge the multisensory time gap: Bimanual coordination to sound and touch with temporal lags. Experimental Brain Research, 235, 135–151. 10.1007/s00221-016-4776-4

Schachner, A., Brady, T. F., Pepperberg, I. M., & Hauser, M. D. (2009). Spontaneous motor entrainment to music in multiple vocal mimicking species. Current Biology, 19, 831–836. 10.1016/j.cub.2009.03.061

Soranzo, A., & Grassi, M. (2014). PSYCHOACOUSTICS: A comprehensive MATLAB toolbox for auditory testing. Frontiers in Psychology, 5, 712. 10.3389/fpsyg.2014.00712

Stiles, L., & Smith, P. F. (2015). The vestibular–basal ganglia connection: Balancing motor control. Brain Research, 1597, 180–188. 10.1016/j.brainres.2014.11.063

Swarbrick, D., Bosnyak, D., Livingstone, S. R., Bansal, J., Marsh-Rollo, S., Woolhouse, M. H., & Trainor, L. J. (2019). How live music moves us: Head movement differences in audiences to live versus recorded music. Frontiers in Psychology, 9, 2682. 10.3389/fpsyg.2018.02682

Takeya, R., Kameda, M., Patel, A. D., & Tanaka, M. (2017). Predictive and tempo-flexible synchronization to a visual metronome in monkeys. Scientific Reports, 7, 6127. 10.1038/s41598-017-06417-3

Tal, I., Large, E. W., Rabinovitch, E., Wei, Y., Schroeder, C. E., Poeppel, D., & Zion Golumbic, E. (2017). Neural entrainment to the beat: The “missing-pulse” phenomenon. Journal of Neuroscience, 37, 6331–6341. 10.1523/jneurosci.2500-16.2017

Tichko, P., Kim, J. C., & Large, E. W. (2021). Bouncing the network: A dynamical systems model of auditory–vestibular interactions underlying infants’ perception of musical rhythm. Developmental Science, 24, e13103. 10.1111/desc.13103

Tichko, P., & Large, E. W. (2019). Modeling infants’ perceptual narrowing to musical rhythms: Neural oscillation and Hebbian plasticity. Annals of the New York Academy of Sciences, 1453, 125–139. 10.1111/nyas.14050

Todd, N. P. M., & Lee, C. S. (2015). The sensory-motor theory of rhythm and beat induction 20 years on: A new synthesis and future perspectives. Frontiers in Human Neuroscience, 9, 444. 10.3389/fnhum.2015.00444

Trainor, L. J., Gao, X., Lei, J.-J., Lehtovaara, K., & Harris, L. R. (2009). The primal role of the vestibular system in determining musical rhythm. Cortex, 45, 35–43. 10.1016/j.cortex.2007.10.014

Truong, D. Q., Thomas, C., Ira, S., Valter, Y., Clark, T. K., & Datta, A. (2024). Unpacking galvanic vestibular stimulation using simulations and relating current flow to reported motions: Comparison across common and specialized electrode placements. PLOS ONE, 19(8), e0309007. 10.1371/journal.pone.0309007

Utz, K. S., Korluss, K., Schmidt, L., Rosenthal, A., Oppenländer, K., Keller, I., & Kerkhoff, G. (2011). Minor adverse effects of galvanic vestibular stimulation in persons with stroke and healthy individuals. Brain Injury, 25, 1058–1069. 10.3109/02699052.2011.607789

Van Dyck, E., Moelants, D., Demey, M., Deweppe, A., Coussement, P., & Leman, M. (2013). The impact of the bass drum on human dance movement. Music Perception, 30, 349–359. 10.1525/mp.2013.30.4.349

Vuust, P., Dietz, M. J., Witek, M., & Kringelbach, M. L. (2018). Now you hear it: A predictive coding model for understanding rhythmic incongruity. Annals of the New York Academy of Sciences, 1423, 19–29. 10.1111/nyas.13622

Vuust, P., Heggli, O. A., Friston, K. J., & Kringelbach, M. L. (2022). Music in the brain. Nature Reviews Neuroscience, 23, 287–305. 10.1038/s41583-022-00578-5

Wang, B., Peterchev, A. V., Gaugain, G., Ilmoniemi, R. J., Grill, W. M., Bikson, M., & Nikolayev, D. (2024). Quasistatic approximation in neuromodulation. Journal of Neural Engineering, 21(4), 041002. 10.1088/1741-2552/ad625e

Wijesinghe, R., Protti, D. A., & Camp, A. J. (2015). Vestibular interactions in the thalamus. Frontiers in Neural Circuits, 9, 79. 10.3389/fncir.2015.00079

Wilkinson, D., Ko, P., Kilduff, P., McGlinchey, R., & Milberg, W. (2005). Improvement of a face perception deficit via subsensory galvanic vestibular stimulation. Journal of the International Neuropsychological Society, 11, 925–929. 10.1017/s1355617705051076

Wing, A. M., Doumas, M., & Welchman, A. E. (2010). Combining multisensory temporal information for movement synchronisation. Experimental Brain Research, 200, 277–282. 10.1007/s00221-009-2134-5

Witek, M. A. G., Clarke, E. F., Kringelbach, M. L., & Vuust, P. (2014). Effects of polyphonic context, instrumentation, and metrical location on syncopation in music. Music Perception, 32, 201–217. 10.1525/mp.2014.32.2.201

Yamazaki, R., & Ushiyama, J. (2024). Head movements induced by voluntary neck flexion stabilize sensorimotor synchronization of the finger to syncopated auditory rhythms. Frontiers in Psychology, 15, 1335050. 10.3389/fpsyg.2024.1335050

Zalta, A., Large, E. W., Schön, D., & Morillon, B. (2024). Neural dynamics of predictive timing and motor engagement in music listening. Science Advances, 10, eadi2525. 10.1126/sciadv.adi2525

